# Unsupervised generative and graph representation learning for modelling cell differentiation

**DOI:** 10.1101/801605

**Authors:** Ioana Bica, Helena Andrés-Terré, Ana Cvejic, Pietro Liò

## Abstract

Using machine learning techniques to build representations from biomedical data can help us understand the latent biological mechanism of action and lead to important discoveries. Recent developments in single-cell RNA-sequencing protocols have allowed measuring gene expression for individual cells in a population, thus opening up the possibility of finding answers to biomedical questions about cell differentiation. In this paper, we explore unsupervised generative neural methods, based on the variational autoencoder, that can model cell differentiation by building meaningful representations from the high dimensional and complex gene expression data. We use disentanglement methods based on information theory to improve the data representation and achieve better separation of the biological factors of variation in the gene expression data. In addition, we use a graph autoencoder consisting of graph convolutional layers to predict relationships between single-cells. Based on these models, we develop a computational framework that consists of methods for identifying the cell types in the dataset, finding driver genes for the differentiation process and obtaining a better understanding of relationships between cells. We illustrate our methods on datasets from multiple species and also from different sequencing technologies.

## Introduction

As technology continues to drive biomedical research forward, new challenges arise with the surge of high volume, information-dense and multivariate data that are generated. The extraction of critical information from such data remains an open problem in biomedical research, which can be significantly aided by the incorporation of machine learning techniques. In particular, unsupervised learning methods have the potential to uncover the underlying structure in biomedical data and therefore propel research on biological processes and diseases that have not yet been fully understood.

This paper aims to build unsupervised neural methods that can be applied to understand cell differentiation using gene expression data. Recent technology for performing single-cell RNA sequencing has resulted in high-throughput experiments capable of measuring gene expression levels for individual cells in a population, thus achieving a granularity not previously possible. In-depth analysis of this high-dimensional and complex gene expression data about the cells can lead to important biomedical discoveries about the factors influencing the differentiation process. However, gene expression data is in general high dimensional, as there are thousands of gene expression measurements for each cell, and very complex.

We propose using a disentangled generative probabilistic framework to model single-cell RNA sequencing gene expression data and build a low dimensional representation that can help us discovering latent biological mechanisms. In this framework, we develop novel methodology that can be used to identify the different cell types in such single-cell RNA-seq datasets using the learned latent representations. We show that we can correctly identify the different cell types in two datasets: one dataset consisting of hematopoietic stem and differentiated cells in zebrafish obtained using Smart-Seq2 [1], and another dataset consisting of humans pancreatic cells obtained using CEL-Seq2 [2]. We also show and discuss some limitations of our methods on a dataset with human hematopoietic cells [3]. In addition, we explore performing perturbations to the latent representation to study how the stem or progenitor cells can be changed into differentiated cells. Moreover, we propose a graph representation learning method based on an autoencoder consisting of graph convolutional layers that can be used to analyze links between single cells.

Current methods for analysing single-cell RNA-seq data are based on the combination of dimensionality reduction techniques and clustering algorithms, at either gene or cell level analysis. The identification of cell lineages and trajectories is one of the main fields of study where single-cell scRNA-seq has had a great influence. The most widespread computational tools include Waterfall or Wishbone [4, 5], which are based on principal component analysis (PCA). Monocle uses independent component analysis (ICA) and SCUBA pseudotime focuses on t-distributed stochastic neighbour embedding (tSNE) [6, 7]. However, some of these methods, particularly the ones based on linear approaches such as PCA, are not able to capture the complex relationships between the input dimensions and can disregard meaningful information within the data. In addition, Yeung and Ruzzo [8] showed that using PCA before clustering gene expression data has a negative effect on the quality of the clusters. Despite these findings, a lot of research in gene expression analysis [9–11] is based on applying PCA before clustering cells to identify their types.

Autoencoders can be used to perform non-linear dimensionality reduction, but also to extract biologically relevant latent features from transcriptomics data. Related work shows the effectiveness of these models in analysing gene expression data. Way *et al*. [12] trained a variational autoencoder on pan-cancer RNA-seq data from The Cancer Genome Atlas [13] to explore the biological relevance of the latent space produced by the autoencoder. Tan *et al*. [14] built a denoising autoencoder capable of modelling the response of cells to low oxygen and finding differences between strains in gene expression from Pseudomonas aeruginosa. Eraslan et al. [15] used autoencoders for denoising purposes, developing a method that is linearly scalable with the number of cells and outperforms existing methods for data imputation. Finally, Rashid et al. [16] used a variational autoencoder to identify tumour subpopulations, marker genes, as well as differentiation trajectories for the malignant cells using scRNA-seq genomic data.

This work represents the first application, to the best of our knowledge, of disentanglement, perturbation and graph-based methods for variational autoencoders with the aim of analysing cell differentiation using single-cell RNA-seq data. We emphasise the importance of building interpretable models, by analysing the relationship between the embedding and gene expression spaces. We also explore the robustness and variability of the latent space by introducing perturbations. Graph representation learning represents a new powerful generation of methodologies for graphs. We show how predicting links between cells can provide insights into differentiation trajectories.

### Disentangled generative probabilistic framework

We propose using a generative probabilistic framework [17] to model the biological processes that lead to the changes in the observed gene expression for cells at different stages in the differentiation process. Let 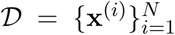 be a high-dimensional single-cell RNAseq dataset consisting of the gene expression of *N* i.i.d cells. Each gene expression vector **x**^(*i*)^ is an observation from a continuous random variable **x**, having distribution *p*_data_(**x**). The gene expression data is assumed to be generated by some random process, modelled by an unobserved continuous random variable **z** with parametrised prior distribution *p*_***θ***_(**z**). The marginal likelihood *p*_***θ***_(x), also known as the evidence, is computed by integrating over the possible latent representations:

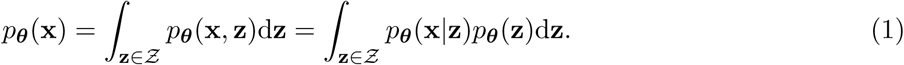

Computing the integral involves spanning the space of values for **z** which is often intractable. For inference, the posterior *p*_***θ***_(**z**|**x**) = (*p*_***θ***_(**x**|**z**)*p*_***θ***_(**z**))*/p*_***θ***_(**x**) has to be computed, which is also intractable, as it requires the marginal likelihood.

To learn in such a framework we use variational inference and we approximate the posterior using the variational distribution *q*_*ϕ*_(**z**|**x**). We thus build a variational autoencoder model [17] and we use a multivariate Gaussian 𝒩 (**z**; ***µ***, diag(***σ***^2^)) distribution with mean ***µ*** and variance ***σ***^2^ to approximate **q**_*ϕ*_(**z**|**x**). An encoder neural network is trained to estimate *q*_***ϕ***_(**z**|**x**). In addition, a isotropic multivariate Gaussian prior is assigned to the latent representation: *p*_***θ***_(**z**) = 𝒩 (**z**; **0, I**). The decoder neural network is trained to reconstruct (generate) the gene expression data from the latent representation and thus estimate *p*_***θ***_(**x**|**z**). See Figure 1.a. for a graphical illustration of the model. The training objective of the standard variational autoencoder model [17] penalises the mutual information between the input gene expression and the latent representation [18] and it also does not encourage disentaglement in the latent representation [19]. Disentanglement is desirable in our case because, ideally, the latent representation **Z** should be able to separate the biological factors that have led to the development of various cell types.

**Figure 1:**
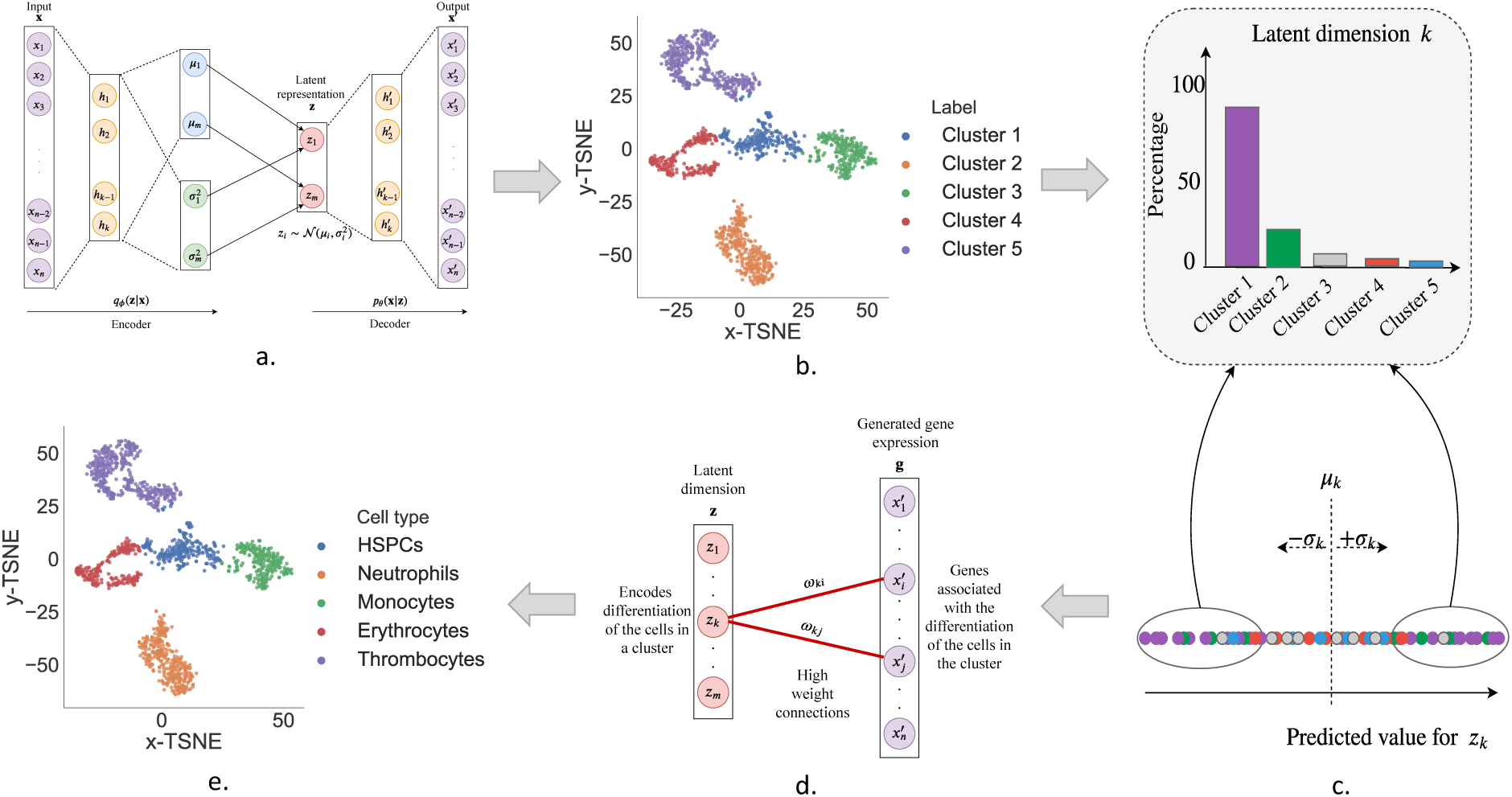
Pipeline for identifying the cell types in a dataset using DiffVAE. Illustration on the zebrafish dataset. **a**. Train DiffVAE to map the gene expression measurements for each cell to a *m*-dimensional latent representation **z. b**. Apply T-SNE on the latent representation **z** and clustering to find the different cell clusters in the dataset. **c**. Identify which latent dimensions in **z** encode the differentiation of the cells in each cluster. **d**. Find the high weights genes for the relevant latent dimensions. **e**. Map the clusters to cell types based on the high weight genes for each cluster.

We introduce DiffVAE, a variational autoencoder that can be used to model and study the differentiation of cells using gene expression data. DiffVAE is part of the MMD-VAE family of autoencoders [19] and is trained to maximize the following objective:

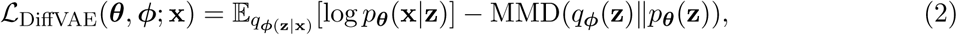

where 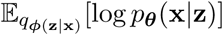 represents the reconstruction accuracy and the maximum mean discrepancy (MMD) [20–22] divergence between *q*_***ϕ***_(**z**) and *p*_***θ***_(**z**) measures how different the moments of two probability distributions are. The intuition behind the MMD divergence is given by the fact that two probability distributions are identical if and only if their moments match. Zhao *et al*. [19] prove that using this training objective will always prefer to maximizes mutual information between the input and the latent representation. Moreover, minimising the divergence MMD(*q*_***ϕ***_(**z**)‖*p*_***θ***_(**z**)), will encourage *q*_***ϕ***_ to be similar to the prior *p*_***θ***_(**z**) = 𝒩 (**z**; **0, I**) with diagonal covariance matrix, which will lead to disentanglement in the latent dimension.

The DiffVAE model consists of two fully connected layers in the decoder and encoder networks. See the Methods section for more details about the DiffVAE model. The methods in this paper were implemented in Python using Keras [23].

### Identifying cell types using DiffVAE

#### Data details and pre-processing

The unsupervised models and the methodology developed in this paper are used to analyse single-cell gene expression data from hematopoietic stem and differentiated cells in zebrafish [1] and in human [3] and also from human pancreatic cells [2]. Let a scRNA-seq dataset be denoted as 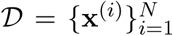, where 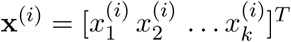 consists of the transcriptomics data for cell (*i*). The zebrafish dataset consists of *k* = 1845 gene expression measurements from *N* = 1422 cells. We used the same 1845 genes identified by [1] to be the most highly variable ones among the 1422 zebrafish single cells. The dataset with human pancreatic cells consists of *N* = 2285 cells with measurements from the *k* = 4000 most highly variable genes. The dataset with human hematopoietic cells contains *N* = 1034 cells with *k* = 700 measurements from the most variable genes. In all cases, we consider that the cell states are initially unknown and we show how the methodology developed in this paper can be used to identify them. Note that some of the results on these datasets are also presented in the supplementary materials.

The transcriptomics data used is log-normalized. However, to use the transcriptomics data as input to DiffVAE, we performed additional normalization through Min-Max scaling such that the expression values for the genes were scaled to the range [0, 1]. This way we model the gene expression for each cell as a multivariate Bernoulli distribution in our probabilistic framework.

#### Pipeline for identifying the cell types

In this section, we describe how DiffVAE can be used to find the different cell types in each dataset. Figure 1 shows the methodological pipeline for this process, with the specific details described in further subsections.

##### USING DIFFVAE TO OBTAIN CELL CLUSTERS

DiffVAE was trained to map the gene expression data for the single cells to a latent representation of *m* dimensions (Figure 1.a). For the datasets with hematopoietic cells (both zebrafish and human), we used *m* = 50 latent dimensions, while for the dataset with the human pancreatic cells, we used *m* = 100 latent dimensions. The large number of latent dimensions is needed to capture the complex biological processes influencing cell differentiation. To visualize the data and identify the different cells, we further use t-Distributed Stochastic Neighbour Embedding (t-SNE) [24] to obtain a 2-dimensional embedding for each cell. In the zebrafish dataset, *K*-means clustering is applied to the t-SNE embedding to obtain 5 cell clusters (Figure 1.b). In the datasets with human cells, we used DBSCAN clustering. We further develop the methodology for mapping each cluster to a cell type.

##### LATENT DIMENSIONS ENCODING CELL DIFFERENTIATION

DiffVAE was designed to model the data generating process giving rise to the observations in our dataset 𝒟. Thus, this method should be able to identify the biological mechanisms that result in the observed gene expression value for our cells. Consider the analysis of a latent dimension *k* for any of the models. Let 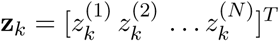 be the predicted value of the encoder for *z*_*k*_ across all of the cells in the dataset. Let *µ*_*k*_ and *σ*_*k*_ be the mean and standard deviation of **z**_*k*_. We define:

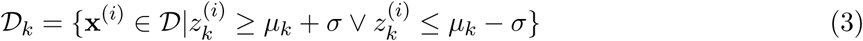

as the set of cells at least a standard deviation from the mean in latent dimension *k*. By computing the percentage distribution of the cells in 𝒟_*k*_ across the distinct cell clusters found in the dataset, we can evaluate how well the latent dimension is encoding the differentiation of the cells in a particular cluster (Figure 1.c). Thus, for each cluster *C* we compute the percentage of cells from cluster *C* in each of 𝒟_*k*_, where *k* ∈ {1, 2,…, 50}. The latent dimensions relevant for the differentiation of cells in cluster *C* will be the ones with the top 10 highest percentage of cells from cluster *C* in 𝒟_*k*_.

##### IDENTIFYING HIGH WEIGHT GENES

The decoder in DiffVAE learns to reconstruct the original gene expression data, and therefore, the weights in the decoder indicate the contribution of each gene in the biological process. By finding the high weight connections between the latent dimensions relevant for each cell cluster and the reconstructed gene expression, we can identify the marker genes for each cell cluster. This will help us identify the cell types.

The high weight connections can be obtained using the weight matrices in the decoder. The decoder consists of a two fully connected layers. Let 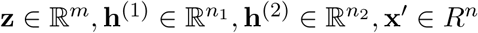, be the sequence of layer activations in the decoder, where the latent dimension **z** represents the input, **h**^(1)^, **h**^(2)^ are the hidden layers and **x**′ is the output. The weight matrices for the connections between the layers in the decoder can be described by 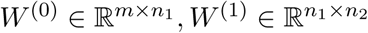. Let ***ω*** ∈ ℝ^*m*×*n*^ be the weight matrix for the connections between the latent dimension and the output. ***ω*** can be computed by multiplying the weight matrices between the individual fully connected layers, as follows: ***ω*** = **W**^(0)^ · **W**^(1)^, where the matrix element *ω*_*ij*_ indicates the weight of the connection between latent dimension *i* and gene *j*. For each latent dimension, the genes are sorted by the absolute value of their weight. The genes having the highest of such weights are referred to the high weight genes. (Figure 1.d)

For each cluster, we selected the latent dimensions that distinguished the best the cells in the clusters and then computed the high weight genes. The high weight genes found for the clusters in the zebrafish dataset are given in Table 1. Using knowledge from biomedical literature about marker genes for blood cells, we mapped each cluster to a cell type. Thus, Cluster 1 corresponds to HSPCs, Cluster 2 to Neutrophils, Cluster 3 to Monocytes, Cluster 4 to Erythrocytes and Cluster 5 to Thrombocytes. The same process was used to map the clusters to cell types in the dataset with human pancreatic cells; see Supplementary Table 1 for the high weight genes found for the clusters in the human pancreatic dataset and their mapping to cell types.

**Table 1:**
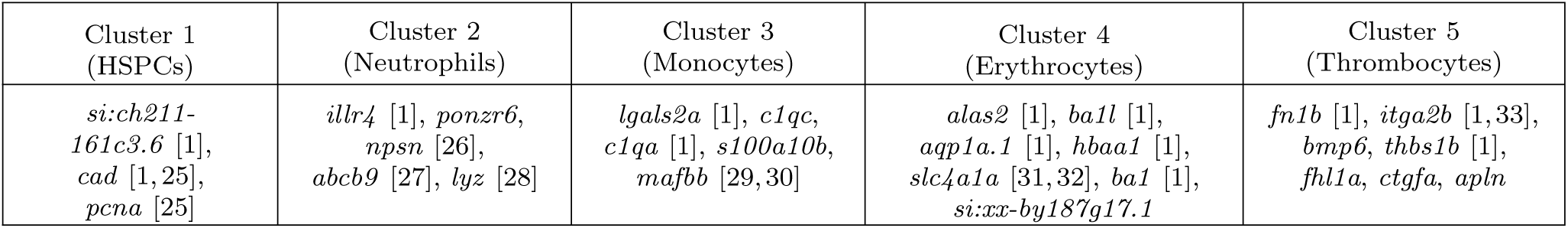
Zebrafish. High weight genes computed using the high weight connections to the latent dimensions with the highest percentage for differentiating the corresponding cell type. Using references from scientific literature each cluster found using DiffVAE is mapped to a cell type.

Our results for identifying the different cell types in the zebrafish dataset are validated by [1] who computationally reconstructed the differentiation trajectories using the Monocle2 algorithm [34] and found the same cellular states. In particular, there is 89.9% overlap between the cell types identified using DiffVAE and the cell types obtained by [1]. Conversely, for the dataset with the human pancreatic cells, we found there is a 96.2% overlap between the cells types obtained using DiffVAE and the ones reported by Murano et al. [2]. In addition, DiffVAE identified all the different cell types in the dataset except for the epsilon cells. However, note that are only 4 epsilon cells in the dataset and Murano et al. [2] also did not identify them computationally, but rather based on the expression of the GHRL gene. See Supplementary Figure 1 for the clusters found using DiffVAE on the dataset with human pancreatic cells.

The representations built by DiffVAE on the human hematopoietic cells do not display separable clusters, which makes it difficult to identify all of the cell types. Velten et al. [3] also indicate that the hematopoietic stem cells, multipotent progenitors and multilymphoid progenitors cells form a unique continuous group when applying clustering methods to the dataset. Refer to Supplementary Figure 2 and the corresponding section for a discussion of the limitations of DiffVAE in this case and directions for future work.

#### Characterization of cell states

For the zebrafish dataset, we also explored the possibility of changing the state of cells through perturbations on the latent dimension. This could help us learn more about the type of biological changes in gene expression that cause a less specialised cell such as an HSPC to differentiate in a more specialised cell such as a Monocyte. For this, we trained a neural network classifier capable of labelling Monocytes, Neutrophils, Erythrocytes and Thrombocytes using the full gene expression data with 99.5% accuracy.

Assume that we have identified, that latent dimension *j* encodes the differentiation of a type of mature blood cells, such as Monocytes. Let *µ*_*j*_ and *σ*_*j*_ be the mean and standard deviation of 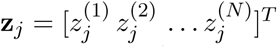 the predicted value of the encoder for *z*_*j*_ across all of the cells in the dataset. We can say that if latent dimension *j* identifies Monocytes, it means that the ratio of the number of Monocytes in 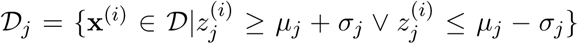 is larger than for the other cells. This strongly suggests that shifting 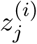 by the standard deviation *σ*_*j*_ of latent dimension *j* could potentially change the cell **x**^(*i*)^ label into a Monocyte.

The method proposed for changing a less specialised cell (an HSPC) into Monocytes involves shifting several of the latent dimensions encoding the differentiation of Monocytes proportionally with their standard deviations. The proportionality factor is the parameter *λ*. The method is illustrated in Figure 3 and it can be generalised to any of the mature cell types that the embedding can separate. Let **x**^(*i*)^ be the input gene expression measurements for cell (*i*). After performing the perturbations on the latent representation **z**^(*i*)^ of cell (*i*) and passing the results through the decoder, we obtain the reconstructed gene expression measurement **y**^(*i*)^. Assume **y**^(*i*)^ is then classified by the neural network as a mature blood cell. By looking at the difference **y**^(*i*)^ − **x**^(*i*)^ one can learn which genes have been more affected by the perturbations on the latent dimension.

**Figure 2:**
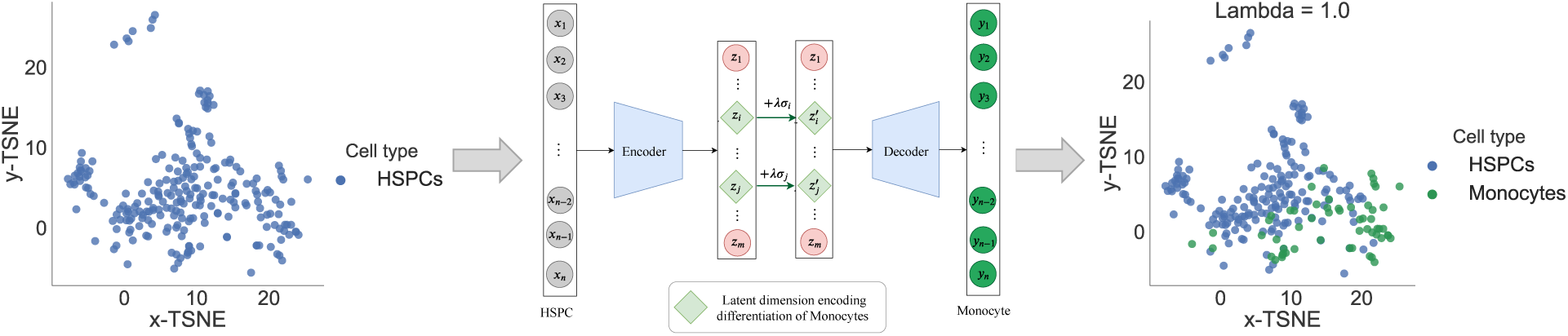
Methodology proposed for changing the cellular states: HSPCs can be converted into Monocytes by shifting the latent dimensions differentiating Monocytes by a factor *λ* multiplied with their standard deviation. Increasing the shifting parameter *λ* will result in more of the HSPCs to be subsequently classified as Monocytes.

**Figure 3:**
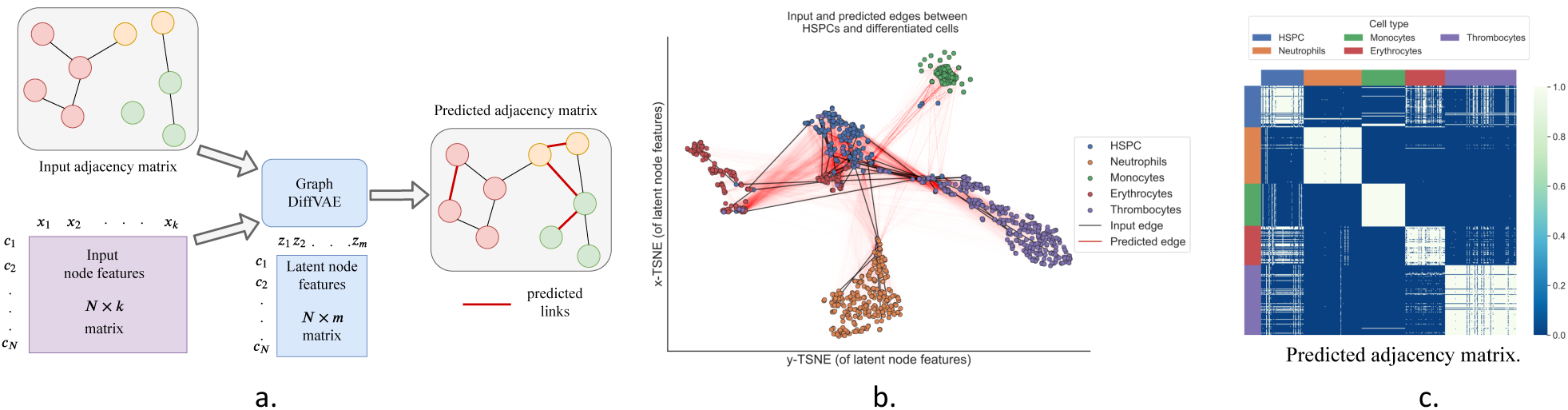
Methodology proposed for analyzing links between cells. **a**. Graph-DiffVAE uses an initial adjacency matrix and individual node features to predict more links between cells. **b**. Projection of cells onto 2-dimensional t-SNE embedding of the latent node features learnt by Graph-DiffVAE and illustration of initial and predicted links between HSPCs and differentiated cells. **c**. Adjacency matrix with predicted links between all cells (the colour white indicates a predicted edge).

#### Comparison of DiffVAE with other dimensionality reduction methods

After performing dimensionality reduction, standard single-cell RNA-seq workflows for identifying cell types involve clustering of the lower representation obtained for the gene expression data [35]. Using the zebrafish cell types found by Athanasiadis et al. [1] and the human pancreatic cell types found by Muraro et al. [2] as true labels, we compare DiffVAE with a standard variational autoencoder (VAE), a simple autoencoder (AE) and Principle Component Analysis (PCA) in terms of clustering performance. Their performance is compared using two clustering algorithms that use different approaches in defining clusters: k-means and DBSCAN. We will cluster both the raw data obtained through dimensionality reduction for *m* ∈ {20, 50, 100} latent dimensions, as well as the 2-dimensional embedding produced using t-SNE.

For each setting of *m* (size of latent dimension), the clustering algorithms (including the computation of the t-SNE embedding) were performed 50 times and each time the ARI between the true labels and the cluster labels was computed. The results reported in Table 2 represent mean ARI obtained on the zebrafish dataset. See Supplementary Table 2 for the results on the dataset with human pancreatic cells. For both datasets, the representation built by DiffVAE gives the best overall clustering performance. In addition, computing the t-SNE embedding on top of the latent representation improves the clustering results.

**Table 2:** Zebrafish. Mean ARI obtained for clustering the latent representation and the t-SNE embedding of the latent representation for three setting of the reduced dimension size *m*. The clustering algorithms used are *k*-means and Gaussian Mixture Models.

| Clustering method | Dim size ($m$) | Latent representation | | | | T-SNE embedding of latent representation | | | |
| --- | --- | --- | --- | --- | --- | --- | --- | --- | --- |
|  |  | DiffVAE | VAE | AE | PCA | DiffVAE | VAE | AE | PCA |
| **k-means** | 20 | **0.803** | 0.771 | 0.799 | 0.633 | **0.809** | 0.738 | 0.699 | 0.717 |
|  | 50 | **0.829** | 0.775 | 0.811 | 0.629 | **0.831** | 0.801 | 0.759 | 0.709 |
|  | 100 | **0.844** | 0.831 | 0.815 | 0.627 | **0.815** | 0.796 | 0.806 | 0.680 |
| **DBSCAN** | 20 | 0.007 | 0.004 | 0.001 | 0.0002 | **0.753** | 0.717 | 0.556 | 0.506 |
|  | 50 | 0.474 | 0.243 | 0.223 | 0.0009 | **0.710** | 0.667 | 0.573 | 0.590 |
|  | 100 | 0.154 | 0.018 | 0.011 | 0.002 | **0.813** | 0.799 | 0.749 | 0.570 |

#### Exploring links between cells

In this section, we shift the focus from just modelling the stochastic behaviour of gene expression across cell types and we also explore modelling the relations between different cell types. For this purpose, we propose Graph-DiffVAE, a graph variational autoencoder where the encoder and the decoder networks are graph convolutional networks. Graph-DiffVAE is based on the graph variational autoencoder proposed by Kipf and Welling [36] and on the Graphite model developed by Grover *et al*. [37].

In this context, we will consider the different cells in the zebrafish dataset as nodes in a graph, represented by the adjacency matrix **A**. The gene expression measurements for each cell will form the node features **X**. The encoder part of Graph-DiffVAE takes as input an initial graph structure for the cells and the input node features and computes a latent representation for cell *q*_***ϕ***_(**Z**|**A, X**), which in this case will be denoted as latent node features. The decoder uses these latent node features and the initial adjacency matrix to predict additional links between the cells, which will be similar to the ones in the input graph.

The input graph can depend on specific applications. One option is to incorporate biological knowledge in the graph, where for instance, edges can represent potential differentiation trajectories for the cells in the dataset. The proposed architecture and training objective for Graph-DiffVAE results in additional edges between cells to be predicted in the output adjacency matrix. The predicted relationships between cells are similar to the ones in the initial graph given as input to the model.

In this paper, we aim to show a proof of concept for using Graph-DiffVAE with single-cell gene expression data. Thus, we propose building an initial graph for the cells where there is an edge between each cell and the cell most similar to it. For this purpose, we will use the Pearson correlation coefficient to measure the similarity between cells. This initial graph is undirected and is represented by a binary adjacency matrix where 1 indicates that there is an edge between two nodes (cells). For each cell in the dataset, we computed the Pearson correlation coefficient between its gene expression vector and the feature vectors of the rest of the cells in the dataset and we added an edge to connect it to the highest positively correlated cell.

Figure 3a. illustrates the pipeline for using Graph-DiffVAE and Figure 3b. shows the 2-dimensional t-SNE embedding of the node features predicted by Graph-DiffVAE, as well as specific links (both initial and predicted) between the HSPCs and differentiated cells, while Figure 3c. show the predicted adjacency matrix of GraphDiff-VAE.

In Figure 3b., it is noticeable that the latent representation built in the encoder exhibits a clustering structure between the different types of cells. This is expected and validates the behaviour of the model, as the cells that are highly correlated to each other are more likely to be part of the same cluster. We can also notice that having an initial edge between these types of cells encourages the prediction of similar types of edges. Moreover, in Figure 4.c this clustering behaviour represented in the encoder is emphasised in the output of the decoder, which predicts relatively well-defined clusters for the Monocytes, Neutrophils, Erythrocytes and Thrombocytes.

An interesting aspect of the predicted graph in Figure 4c. is the fact that the HSPCs do not cluster together that well. In particular, there are clear links between several HSPCs and all of the other cells in a cluster of mature blood cells. This means that among the HSPCs there are cells that have already started the process of differentiation towards one of the specific mature cells. Additionally, we can notice that Graph-DiffVAE predicted more edges between HSPCs and Erythrocytes compared to the other differentiated cells.

## Discussion and conclusion

In this paper, we explored unsupervised generative and graph representation learning methods for modelling single-cell gene expression data and understanding cell differentiation by developing the DiffVAE and Graph-DiffVAE models. The two different models succeed in characterising different states of cell differentiation based on single-cell RNA-Seq data. We illustrated how to identify cell types using DiffVAE through a pipeline that involves clustering the latent representation, detecting important genes for each cluster and mapping from clusters to cells type. We have shown that the pipeline is applicable to datasets of different nature, providing powerful insight into the noisy information concealed by single-cell genetic data.

Moreover, the embeddings created constitute a set of generative functions that can produce artificial samples, allowing further exploration and expansion of the current datasets. Additionally, we explored perturbations over the generative latent space to then analyse the effect on the gene expression and changes in cellular states. That can lead to future studies on the stability of cellular states, and robustness over genetic stochasticity.

Finally, through Graph-DiffVAE we explored a way of understanding the connections between cells, and in particular between HSPCs and differentiated cells. Further analysis in this direction can be useful in quantifying the degree of differentiation or stemness of the HSPCs and studying differentiation trajectories.

From a methodological perspective, future work could involve combining DiffVAE and Graph-DiffVAE into a single multitask learning framework [38] and using Graph Attentional Layers [39] as part of Graph-DiffVAE to better quantifying the importance of the links between cells.

## Methods

### DiffVAE

DiffVAE was constructed such that both the encoder and decoder consists of two fully connected hidden consisting of 512 and 256 neurons respectively. The incorporation of multiple hidden layers helps to build a hierarchical representation of features, thus obtaining a more complex model. The size of the hidden layers is symmetric between the encoder and decoder. The latent representation **z** has *m* = 50 dimensions for the hematopoietic datasets and *m* = 100 dimensions of the dataset with human pancreatic cells. The ReLU activation was applied in the hidden layers of both the encoder and decoder in order to introduce non-linearity in the network. The specific operations performed by DiffVAE are as follows:

### Encoder (Inference model): q_*ϕ*_(z|x) = 𝒩 (z; *µ*, diag(*σ*^2^))

The encoder consists of fully connected layers and has a Gaussian output. For numerical stability, the encoder network learns log(***σ***^2^) instead of ***σ***^2^. The input to the encoder is **x** ∈ ℝ^1×1845^, which, in our case, represents the gene expression data. The operations performed by the encoder network are summarised by:

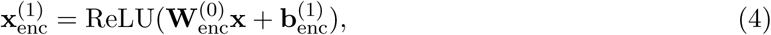

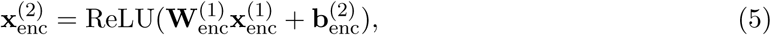

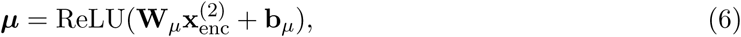

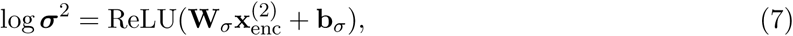

where 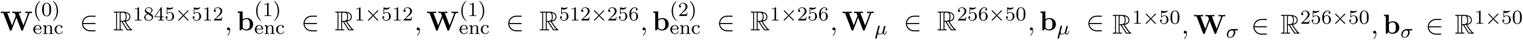 are the trainable parameters in the encoder. The encoder also uses batch normalization [40] to overcome the problem of internal covariate shift.

Directly sampling the latent representation **z** can cause problems to the standard gradient-based algorithm, as it is not possible to compute gradients through the random sampling of **z**. To overcome these issues, Kingma and Welling [17] proposed the reparameterisation trick that involves parameterising the latent code as follows:

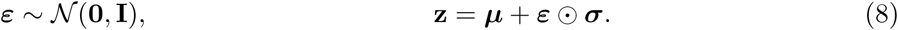

### Decoder (generative model): *p*_*θ*_(x|z)

The output of the decoder has to reward the likelihood of the data we want to generate with this model. In our case, for each data point, the gene expression values can be modelled as samples from a multivariate Bernoulli distribution. Intuitively, each input gene is modelled as a Bernoulli random variable, and a sample from this distribution indicates whether the gene is expressed or not. To build a decoder with Bernoulli output, we need to apply the logistic activation function to compute the output of the decoder because it takes values in the range [0, 1].

The input to the decoder is the latent representation **z**. The decoder performs the following operations in order to obtain the reconstructed input **x**′:

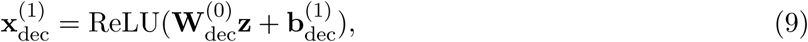

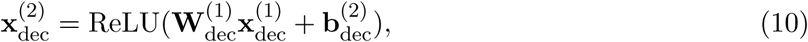

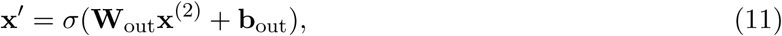

where *σ* is the logistic activation function and 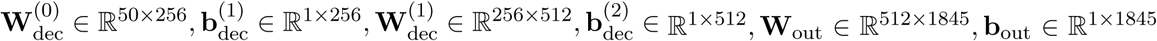 are the trainable parameters in the decoder. As **x**′ is not sampled, we provide a maximum likelihood estimate for the reconstruction.

DiffVAE was trained using minibatch stochastic gradient descent to minimize −ℒ_DiffVAE_:

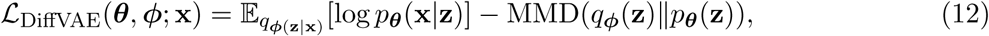

We used the Adam Optimizer with a learning rate of 0.001 and a batch size of 128 and we trained DiffVae for 100 epochs.

### Additional models used as benchmarks

To assess the performance of DiffVAE on clustering, we compare it against the following benchmarks: standard VAE, standard autoencoder and PCA. For the standard VAE and the standard autoencoder, we used the same number of layers and neurons as we used for DiffVAE. For both models, we also used the Adam Optimizer with a learning rate of 0.001, a batch size of 128 and we trained them for 100 epochs.

The objective function maximised by the variational autoencoder is:

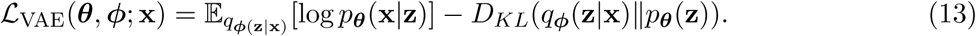

The standard autoencoder model is trained to minimise the reconstruction error, which can be measured by the mean squared error loss function *l*(**x, *ω***) = ‖**x** − *dec*(*enc*(**x**)) ‖^2^, where ***ω*** consists of all of the parameters in the encoder and decoder networks.

The neural network trained for classifying the mature cell types takes as input the gene expression data for the cell and output the probability that the particular cell is a Monocyte, Neutrophil, Thrombocyte or Erythrocyte. The model consists of three hidden layers of sizes 256, 512, 256 neurons with ReLU activation and an output layer with 4 neurons and softmax activation. The model was also trained using the Adam Optimizer for 300 epochs with a learning rate of 0.001 and a batch size of 128.

### GraphDiffVAE

Let 𝒢 = (𝒱, *ε*), with |𝒱| = *N* be an undirected and unweighted initial graph built from the cells, defined by the binary adjacency matrix **A** ∈ {0, 1}^*N*×*N*^. Such an initial graph for the cells can be already available or it can be artificially built using the Pearson correlation between cells.

Let **X** be an *N* × *F* matrix consisting of node features, where *F* is the number of features for each node. In our case, the nodes are the different cells in the dataset, and the features are represented by the gene expression for each cell. Assume that each node is connected to itself, so that **A** has diagonal entries *A*_*ii*_ = 1. Let **D** be the diagonal degree matrix of **A**: *D*_*ii*_ = Σ_*j*_ *A*_*ij*_.

Graph convolutional networks (GCN) were proposed by Kipf and Welling [41] with the following layer wise propagation rule:

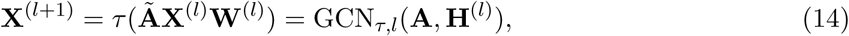

where **Ã** = **D**^−1*/*2^**AD**^−1*/*2^, *τ* is the activation function applied, **X**^(*l*)^ and **X**^(*l*+1)^ are the activations of the layers (*l*) and (*l* + 1) respectively, and **W**^(*l*)^ are the weights. **X**^(0)^ represents the input feature matrix **X**. The size of layer *n*_*l*_, represents by the number of node features computed at layer (*l*).

Through the layer-wise propagation rule, the graph convolutional network performs spectral graph convolutions. The model can be regarded as the differentiable and generalised version of the algorithm proposed by Weisfeiler-Lehman on graphs. In particular, the layer-wise propagation rule can be viewed as a message passing computation over the graph structure. Through one hidden layer, nodes in the graph pass information about their local structure to neighbours that are 1-hop away. Based on the information received from the neighbours, the nodes update their node features.

Graph-DiffVAE is a graph variational autoencoder where the encoder and the decoder networks are graph convolutional networks applying the layer-wise propagation rule. The architecture of Graph-DiffVAE is based on the ones in the graph variational autoencoder proposed by Kipf and Welling [36] and in the Graphite model developed by Grover *et al*. [37].

### Inference model (encoder) : *q*_*ϕ*_(Z|A, X)

The encoder in Graph-DiffVAE is represented by a graph convolutional network with multiple layers and with Gaussian output. The input to the encoder consists of the matrix with node features **X** and of the graph adjacency matrix **A**. The layers in the encoder network perform the following operations:

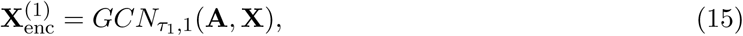

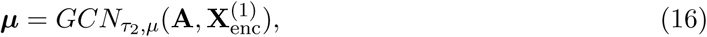

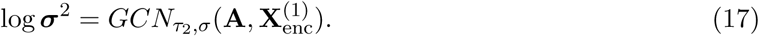

where *τ*_1_ is the ReLU activation function and *τ*_2_ is the linear activation function. The number of node features computed in 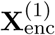 is 512 and the number of node features in ***µ*** and ***σ***^2^ is 50, thus forming a latent representation **Z** where each node has *M* = 50 features.

The encoder represents a factorised multivariate Gaussian distribution, such that:

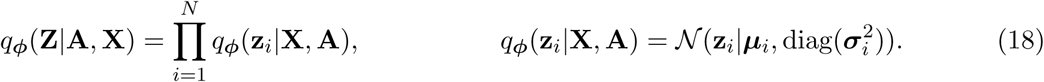

The reparametrisation trick is used again to sample each **z**_*i*_:

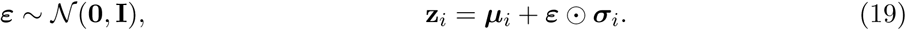

It is important to notice that the latent representation **Z** build by the Graph-DiffVAE encoder contains information from both the graph structure and the node features. **Z** encompass the latent representations for each node in the graph.

### Generative model (decoder): *p*_*ϕ*_(A|Z, X)

The output of the decoder is an adjacency matrix **Â** representing an undirected and unweighted graph with predicted edges between nodes. Such an adjacency matrix can be represented by a factorised Bernoulli distribution.

The decoder network uses as input the initial adjacency matrix **A** and a concatenation of the input node features **X** and the latent node features computed by the encoder **Z**, described by [**Z**|**X**]. The layers in the decoder network perform the following operations:

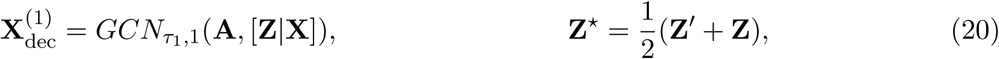

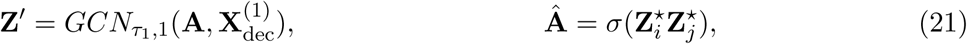

where *τ*_1_ is the ReLU activation function. The number of node features computed in 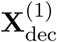 is 512. The decoder builds its own latent representation **Z**′ consisting of 50 node features which it then adds to the representation constructed through the encoder to obtain **Z**^⋆^.

Similarly with the standard framework of the variational autoencoder, we optimize the following objective for Graph-DiffVAE:

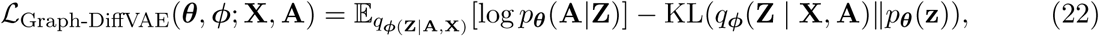

The model was trained for 200 epochs with a learning rate of 0.0001. The proposed architecture and training objective for Graph-DiffVAE results in additional edges between cells to be predicted in the output adjacency matrix **Â**. The predicted relationships between cells are similar to the ones in the initial graph given as input to the model.

### Software

The code for DiffVAE and Graph-DiffVAE is publicly available in the GitHub repository: https://github.com/ioanabica/DiffVAE.

## Data availability

The zebrafish dataset used for this paper is made publicly available by Athanasiadis et al. [1] on ArrayExpress under the accession numbers E-MTAB-3947, E-MTAB-4617 and E-MTAB-5530 and also at https://www.sanger.ac.uk/science/tools/basicz. Similarly, the dataset with human pancreatic cells is made publicly available by Murarot al. [2] under accession number GEO: GSE85241. Moreover, the dataset with human hematopoietic cells was made publicly available by Velten et al. [3] under accession code GEO: GSE75478.

## Acknowledgements

I.B. is funded by The Alan Turing Institute Doctoral Studentship, under the EPSRC grant EP/N510129/1.

## Results on dataset with human pancreatic cells

**Supplementary Figure 1:**
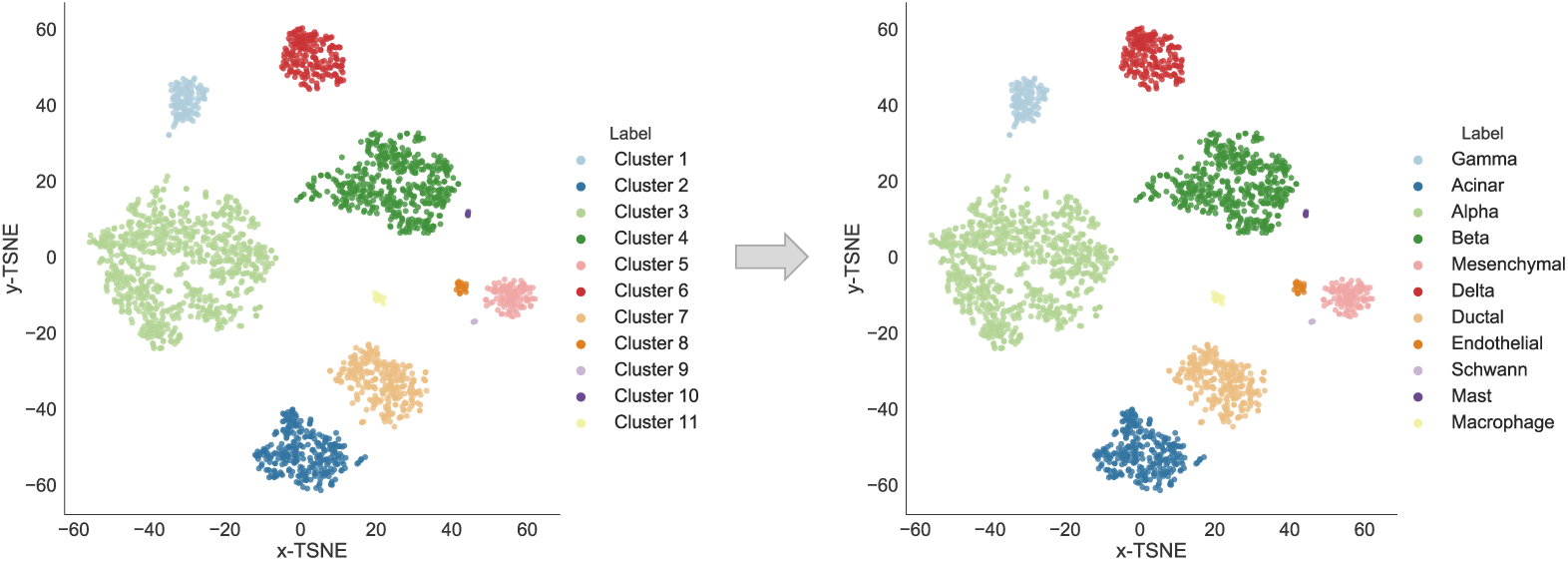
Clusters identified by DiffVAE in the dataset with human pancreatic cells.

**Supplementary Table 1:** High weight genes computed using the high weight connections to the latent dimensions with the highest percentage for differentiating the corresponding cell type. Using references from scientific literature each cluster found using DiffVAE in the dataset with human pancreatic cells is mapped to a cell type.

| Cluster 1<br>(Gamma) | Cluster 2<br>(Acinar) | Cluster 3<br>(Alpha) | Cluster 4<br>(Beta) | Cluster 5<br>(Mesenchymal) | Cluster 6<br>(Delta) |
| --- | --- | --- | --- | --- | --- |
| *PPY*, *SERTM1*,<br>*CARTPT*, *LMO3*,<br>*ZNF503* | *PRSS1* [2],<br>*REG1B* [2],<br>*ALB* [2] | *GCG* [2],<br>*LOXL4* [2],<br>*CRYBA2* [2],<br>*IRX2*, *FEV* [2, 42] | *MAFA* [2],<br>*INS* [2], *SIX2* [2], | *COL1A1* [2],<br>*COL1A2* [2],<br>*COL3A1* [2],<br>*SPARC* [2] | *SST* [2], *RBP4* [2],<br>*GABRG2* [2],<br>*GHSR* [2] |
| Cluster 7<br>(Ductal) | Cluster 8<br>(Endothelial) | Cluster 9<br>(Schwann) | Cluster 10<br>(Mast) | Cluster 11<br>(Macrophage) |  |
| *KRT19* [2],<br>*SPP1* [2], | *SOX18* [2],<br>*SNAI1* [2],<br>*BCL6B* [2] | *NPY*, *GFRA3* [43] | *HDC* [44],<br>*TPSAB1* [45],<br>*KIT* [46, 47] | *C3AR1* [48],<br>*CD300A* [49],<br>*STAB1* [50] |  |

**Supplementary Table 2:** Human pancreatic cells. Mean ARI obtained for clustering the latent representation and the t-SNE embedding of the latent representation for three setting of the reduced dimension size *m*.

| Clustering method | Dim size ($m$) | Latent representation | | | | T-SNE embedding of latent representation | | | |
| --- | --- | --- | --- | --- | --- | --- | --- | --- | --- |
|  |  | DiffVAE | VAE | AE | PCA | DiffVAE | VAE | AE | PCA |
| **k-means** | 20 | **0.678** | 0.605 | 0.527 | 0.549 | 0.683 | 0.689 | **0.697** | 0.689 |
|  | 50 | **0.636** | 0.612 | 0.525 | 0.283 | **0.697** | 0.681 | 0.694 | 0.654 |
|  | 100 | 0.607 | 0.605 | **0.633** | 0.557 | **0.706** | 0.686 | 0.676 | 0.658 |
| **DBSCAN** | 20 | 0.020 | 0.0001 | 0.021 | 0.002 | **0.932** | 0.927 | 0.887 | 0.837 |
|  | 50 | 0.292 | 0.001 | 0.073 | 0.008 | **0.933** | 0.891 | 0.856 | 0.865 |
|  | 100 | 0.345 | 0.021 | 0.0005 | 0.042 | **0.957** | 0.943 | 0.867 | 0.853 |

## Results on dataset with human hematopoietic cells

**Supplementary Figure 2.**
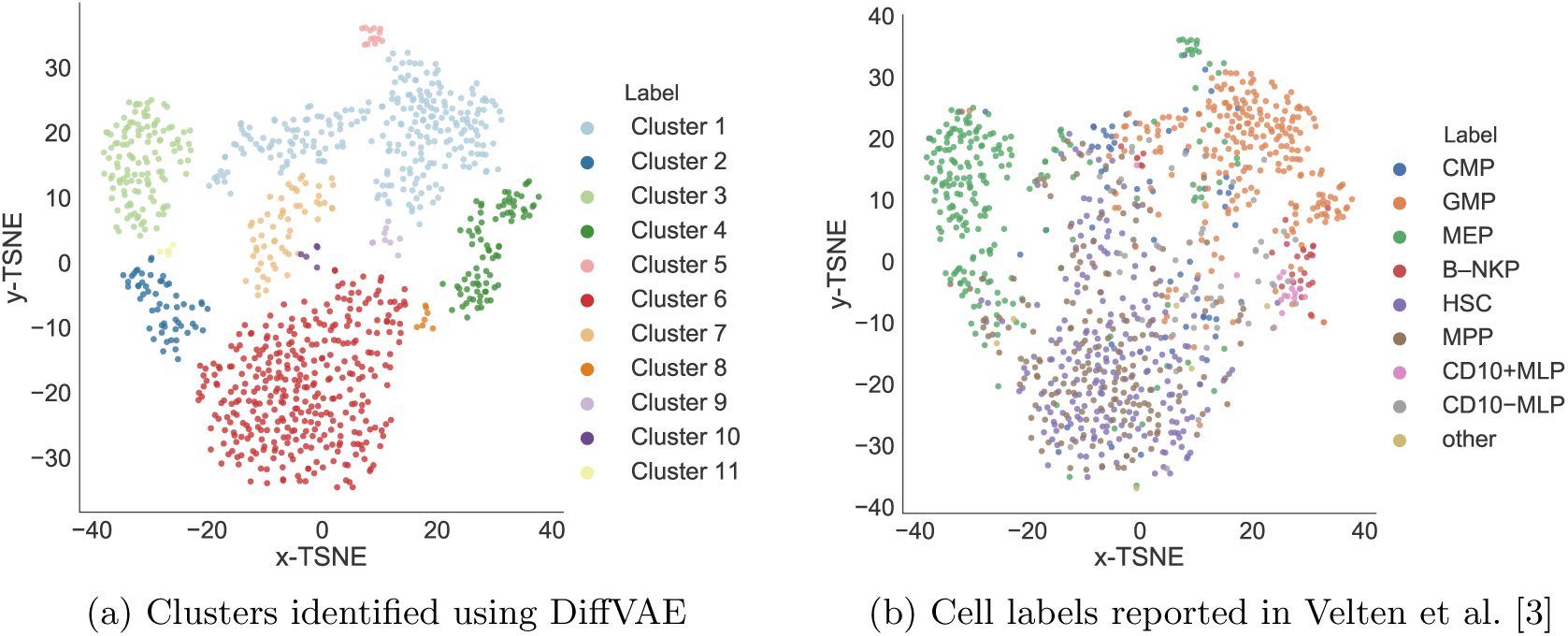

Supplementary Figure 2 As it can be noticed in Supplementary Figure 2, the latent representation obtained through DiffVAE does not produce well-defined cell clusters for the dataset with human hematopoietic cells. Supplementary Figure 2 (a) shows the clusters obtained when using DBSCAN on the T-SNE embedding computed on top of the 50-dimensional representation obtained through DiffVAE. The high weight genes for Cluster 2 are *GATA2, GATA1, RRM2, ITGA2B, MYBL2* and for Cluster 3 are *GATA1, TYMS, KLF1, TFR2* which helps us determine that Clusters 2 and 3 contain the MEP cells. High weight genes for cluster 1 include: *CTSG, AZU1, ELANE, LYZ, ELANE, LGMN* indicates that the GMP cells are part of this cluster. Moreover, the presence of B cells in cluster 4 is indicated by some of the high weight genes for cluster 4: *DNTT, VPREB1, JCHAIN*.

Nevertheless, in Supplementary Figure 2 (b) we plotted the cell labels (based on FACS surface phenotype) provided by Velten et al. [3]. We notice that the HSC and MPP cells cluster together. Similarly, the CMP cells are also scattered and do not form a cluster. Velten et al. [3] reported similar results when applying clustering methods to this dataset.

The limitations of DiffVAE for this dataset may be caused by a large number of different cell states compared to the number of samples available to distinguish between them. This problem is also amplified by the fact that the cell states are close to each other in the process of haematopoiesis.

## References

[1] E. I. Athanasiadis, J. G. Botthof, H. Andres, L. Ferreira, P. Lio, and A. Cvejic, “Single-cell rna-sequencing uncovers transcriptional states and fate decisions in haematopoiesis,” Nature communications, vol. 8, no. 1, p. 2045, 2017.

[2] M. J. Muraro, G. Dharmadhikari, D. Grün, N. Groen, T. Dielen, E. Jansen, L. van Gurp, M. A. Engelse, F. Carlotti, E. J. de Koning, et al., “A single-cell transcriptome atlas of the human pancreas,” Cell systems, vol. 3, no. 4, pp. 385–394, 2016.

[3] L. Velten, S. F. Haas, S. Raffel, S. Blaszkiewicz, S. Islam, B. P. Hennig, C. Hirche, C. Lutz, E. C. Buss, D. Nowak, et al., “Human haematopoietic stem cell lineage commitment is a continuous process,” Nature cell biology, vol. 19, no. 4, p. 271, 2017.

[4] e. a. Shin, Jaehoon, “Single-cell rna-seq with waterfall reveals molecular cascades underlying adult neurogenesis,” Cell stem cell, vol. 17, no. 3, pp. 360–372, 2015.

[5] e. a. Setty, Manu, “Wishbone identifies bifurcating developmental trajectories from single-cell data.,” Nature biotechnology, vol. 34, no. 6, p. 637, 2016.

[6] e. a. Trapnell, Cole, “The dynamics and regulators of cell fate decisions are revealed by pseudotemporal ordering of single cells.,” Nature biotechnology, vol. 32, no. 4, p. 381, 2014.

[7] e. a. Marco, Eugenio, “Bifurcation analysis of single-cell gene expression data reveals epigenetic landscape.,” Proceedings of the National Academy of Sciences, vol. 111, no. 52, pp. E5643–E5650, 2014.

[8] K. Y. Yeung and W. L. Ruzzo, “Principal component analysis for clustering gene expression data,” Bioinformatics, vol. 17, no. 9, pp. 763–774, 2001.

[9] C. Guibentif, R. E. Rönn, C. Böiers, S. Lang, S. Saxena, S. Soneji, T. Enver, G. Karlsson, and N.-B. Woods, “Single-cell analysis identifies distinct stages of human endothelial-to-hematopoietic transition,” Cell reports, vol. 19, no. 1, pp. 10–19, 2017.

[10] S. McKinney-Freeman, P. Cahan, H. Li, S. A. Lacadie, H.-T. Huang, M. Curran, S. Loewer, O. Naveiras, K. L. Kathrein, M. Konantz, et al., “The transcriptional landscape of hematopoietic stem cell ontogeny,” Cell stem cell, vol. 11, no. 5, pp. 701–714, 2012.

[11] Y. Kluger, D. P. Tuck, J. T. Chang, Y. Nakayama, R. Poddar, N. Kohya, Z. Lian, A. B. Nasr, H. R. Halaban, D. S. Krause, et al., “Lineage specificity of gene expression patterns,” Proceedings of the National Academy of Sciences of the United States of America, vol. 101, no. 17, pp. 6508–6513, 2004.

[12] G. P. Way and C. S. Greene, “Extracting a biologically relevant latent space from cancer transcriptomes with variational autoencoders,” bioRxiv, p. 174474, 2017.

[13] J. N. Weinstein, E. A. Collisson, G. B. Mills, K. R. M. Shaw, B. A. Ozenberger, K. Ellrott, I. Shmulevich, C. Sander, J. M. Stuart, C. G. A. R. Network, et al., “The cancer genome atlas pan-cancer analysis project,” Nature genetics, vol. 45, no. 10, p. 1113, 2013.

[14] J. Tan, J. H. Hammond, D. A. Hogan, and C. S. Greene, “Adage-based integration of publicly available pseudomonas aeruginosa gene expression data with denoising autoencoders illuminates microbe-host interactions,” MSystems, vol. 1, no. 1, pp. e00025–15, 2016.

[15] e. a. Eraslan, Gkcen, “Single-cell rna-seq denoising using a deep count autoencoder.,” Nature communications, vol. 10, no. 1, p. 390, 2019.

[16] S. Rashid, S. Shah, Z. Bar-Joseph, and R. Pandya, “Project dhaka: Variational autoencoder for unmasking tumor heterogeneity from single cell genomic data,” bioRxiv, p. 183863, 2018.

[17] D. P. Kingma and M. Welling, “Auto-encoding variational bayes,” arXiv preprint 1312.6114, 2013.

[18] N. Tishby and N. Zaslavsky, “Deep learning and the information bottleneck principle,” in Information Theory Workshop (ITW), 2015 IEEE, pp. 1–5, IEEE, 2015.

[19] S. Zhao, J. Song, and S. Ermon, “Infovae: Information maximizing variational autoencoders,” arXiv preprint 1706.02262, 2017.

[20] A. Gretton, K. M. Borgwardt, M. Rasch, B. Schölkopf, and A. J. Smola, “A kernel method for the two-sample-problem,” in Advances in neural information processing systems, pp. 513–520, 2007.

[21] Y. Li, K. Swersky, and R. Zemel, “Generative moment matching networks,” in International Conference on Machine Learning, pp. 1718–1727, 2015.

[22] G. K. Dziugaite, D. M. Roy, and Z. Ghahramani, “Training generative neural networks via maximum mean discrepancy optimization,” arXiv preprint 1505.03906, 2015.

[23] F. Chollet et al., “Keras,” 2015.

[24] L. v. d. Maaten and G. Hinton, “Visualizing data using t-sne,” Journal of machine learning research, vol. 9, no. Nov, pp. 2579–2605, 2008.

[25] A. Y. Leung, J. C. Leung, L. Y. Chan, E. S. Ma, T. T. Kwan, K. Lai, A. Meng, and R. Liang, “Proliferating cell nuclear antigen (pcna) as a proliferative marker during embryonic and adult zebrafish hematopoiesis,” Histochemistry and cell biology, vol. 124, no. 2, pp. 105–111, 2005.

[26] P. Patil, T. Uechi, and N. Kenmochi, “Incomplete splicing of neutrophil-specific genes affects neutrophil development in a zebrafish model of poikiloderma with neutropenia,” RNA biology, vol. 12, no. 4, pp. 426–434, 2015.

[27] M. J. Foulkes, K. M. Henry, J. Rougeot, E. Hooper-Greenhill, C. A. Loynes, P. Jeffrey, A. Fleming, C. O. Savage, A. H. Meijer, S. Jones, et al., “Expression and regulation of drug transporters in vertebrate neutrophils,” Scientific reports, vol. 7, no. 1, p. 4967, 2017.

[28] E. A. Harvie and A. Huttenlocher, “Neutrophils in host defense: new insights from zebrafish,” Journal of leukocyte biology, vol. 98, no. 4, pp. 523–537, 2015.

[29] M. T. N. Tran, M. Hamada, H. Jeon, R. Shiraishi, K. Asano, M. Hattori, M. Nakamura, Y. Imamura, Y. Tsunakawa, R. Fujii, et al., “Mafb is a critical regulator of complement component c1q,” Nature communications, vol. 8, no. 1, p. 1700, 2017.

[30] L. M. Kelly, U. Englmeier, I. Lafon, M. H. Sieweke, and T. Graf, “Mafb is an inducer of monocytic differentiation,” The EMBO journal, vol. 19, no. 9, pp. 1987–1997, 2000.

[31] W. Pimtong, M. Datta, A. M. Ulrich, and J. Rhodes, “Drl. 3 governs primitive hematopoiesis in zebrafish,” Scientific reports, vol. 4, p. 5791, 2014.

[32] F. E. Moore, E. G. Garcia, R. Lobbardi, E. Jain, Q. Tang, J. C. Moore, M. Cortes, A. Molodtsov, M. Kasheta, C. C. Luo, et al., “Single-cell transcriptional analysis of normal, aberrant, and malignant hematopoiesis in zebrafish,” Journal of Experimental Medicine, pp. jem–20152013, 2016.

[33] G. Khandekar, S. Kim, and P. Jagadeeswaran, “Zebrafish thrombocytes: functions and origins,” Advances in hematology, vol. 2012, 2012.

[34] X. Qiu, A. Hill, J. Packer, D. Lin, Y.-A. Ma, and C. Trapnell, “Single-cell mrna quantification and differential analysis with census,” Nature methods, vol. 14, no. 3, p. 309, 2017.

[35] M. D. Luecken and F. J. Theis, “Current best practices in single-cell rna-seq analysis: a tutorial,” Molecular systems biology, vol. 15, no. 6, 2019.

[36] T. N. Kipf and M. Welling, “Variational graph auto-encoders,” arXiv preprint 1611.07308, 2016.

[37] A. Grover, A. Zweig, and S. Ermon, “Graphite: Iterative generative modeling of graphs,” arXiv preprint 1803.10459, 2018.

[38] Y. Zhang and Q. Yang, “A survey on multi-task learning,” arXiv preprint 1707.08114, 2017.

[39] P. Veličković, G. Cucurull, A. Casanova, A. Romero, P. Liò, and Y. Bengio, “Graph attention networks,” arXiv preprint 1710.10903, 2017.

[40] S. Ioffe and C. Szegedy, “Batch normalization: Accelerating deep network training by reducing internal covariate shift,” arXiv preprint 1502.03167, 2015.

[41] T. N. Kipf and M. Welling, “Semi-supervised classification with graph convolutional networks,” arXiv preprint 1609.02907, 2016.

[42] M. E. Kimple, M. P. Keller, M. R. Rabaglia, R. L. Pasker, J. C. Neuman, N. A. Truchan, H. K. Brar, and A. D. Attie, “Prostaglandin e2 receptor, ep3, is induced in diabetic islets and negatively regulates glucose-and hormone-stimulated insulin secretion,” Diabetes, vol. 62, no. 6, pp. 1904–1912, 2013.

[43] J. Widenfalk, A. Tomac, E. Lindqvist, B. Hoffer, and L. Olson, “Gfr*α*-3, a protein related to gfr*α*-1, is expressed in developing peripheral neurons and ensheathing cells,” European journal of neuroscience, vol. 10, no. 4, pp. 1508–1517, 1998.

[44] A. Kuramasu, H. Saito, S. Suzuki, T. Watanabe, and H. Ohtsu, “Mast cell-/basophil-specific transcriptional regulation of human l-histidine decarboxylase gene by cpg methylation in the promoter region,” Journal of Biological Chemistry, vol. 273, no. 47, pp. 31607–31614, 1998.

[45] G. H. Caughey, “Mast cell tryptases and chymases in inflammation and host defense,” Immunological reviews, vol. 217, no. 1, pp. 141–154, 2007.

[46] A. C. Garcia-Montero, M. Jara-Acevedo, C. Teodosio, M. L. Sanchez, R. Nunez, A. Prados, I. Aldanondo, L. Sanchez, M. Dominguez, L. M. Botana, et al., “Kit mutation in mast cells and other bone marrow hematopoietic cell lineages in systemic mast cell disorders: a prospective study of the spanish network on mastocytosis (rema) in a series of 113 patients,” Blood, vol. 108, no. 7, pp. 2366–2372, 2006.

[47] G. Cruse, D. D. Metcalfe, and A. Olivera, “Functional deregulation of kit: link to mast cell proliferative diseases and other neoplasms,” Immunology and Allergy Clinics, vol. 34, no. 2, pp. 219–237, 2014.

[48] Y. Mamane, C. C. Chan, G. Lavallee, N. Morin, L.-J. Xu, J. Huang, R. Gordon, W. Thomas, J. Lamb, E. E. Schadt, et al., “The c3a anaphylatoxin receptor is a key mediator of insulin resistance and functions by modulating adipose tissue macrophage infiltration and activation,” Diabetes, vol. 58, no. 9, pp. 2006–2017, 2009.

[49] O. Zenarruzabeitia, J. Vitallé, C. Eguizabal, V. R. Simhadri, and F. Borrego, “The biology and disease relevance of cd300a, an inhibitory receptor for phosphatidylserine and phosphatidylethanolamine,” The Journal of Immunology, vol. 194, no. 11, pp. 5053–5060, 2015.

[50] P. Rantakari, D. A. Patten, J. Valtonen, M. Karikoski, H. Gerke, H. Dawes, J. Laurila, S. Ohlmeier, K. Elima, S. G. Hübscher, et al., “Stabilin-1 expression defines a subset of macrophages that mediate tissue homeostasis and prevent fibrosis in chronic liver injury,” Proceedings of the National Academy of Sciences, vol. 113, no. 33, pp. 9298–9303, 2016.

